## Supplemental Figures for "Immortalization of different breast epithelial cell types results in distinct mitochondrial mutagenesis"

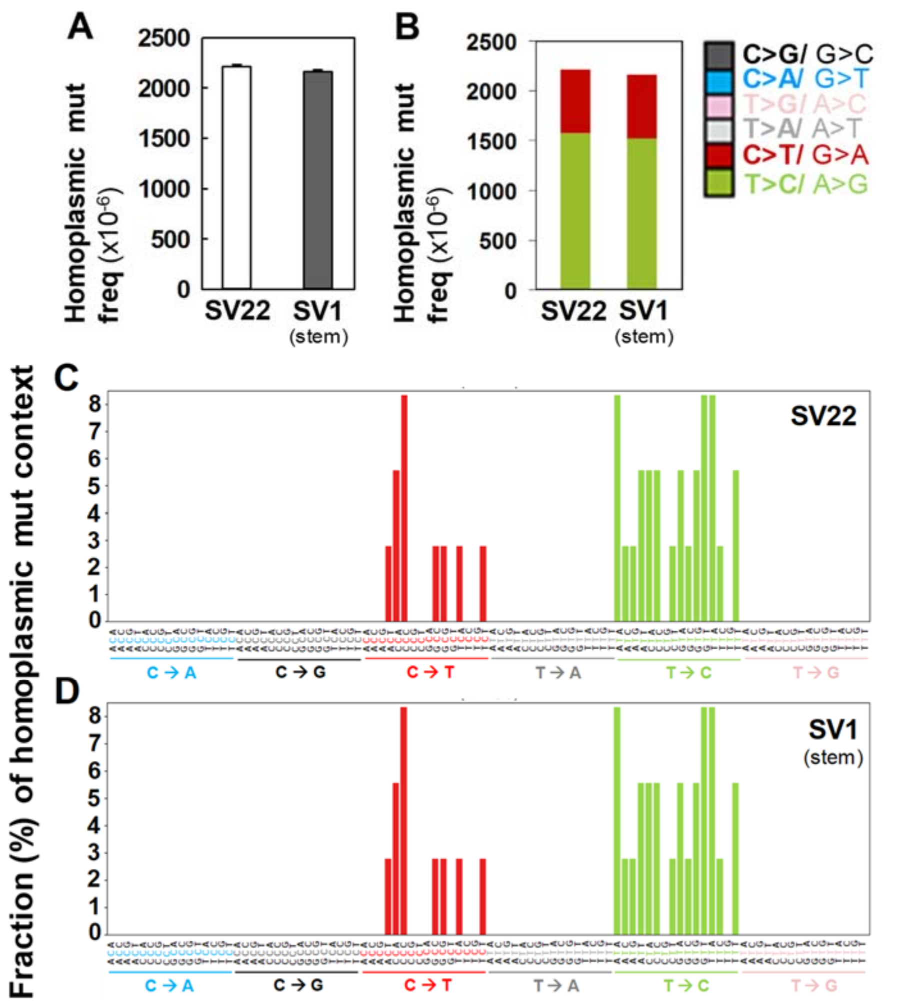


**Figure S1. Frequencies and fractions (%) of sequence context spectra of homoplasmic mutations in the whole mtDNA.** Overall homoplasmic mutation frequency (**A**), frequencies of each type of homoplasmic mutation (**B**), and homoplasmic unique mutation context spectra (**C**,**D**) for SV22 (immortalized non-stem) and SV1 (immortalized stem) cells were determined using Duplex Sequencing. Error bars represent the Wilson Score 95% confidence intervals.

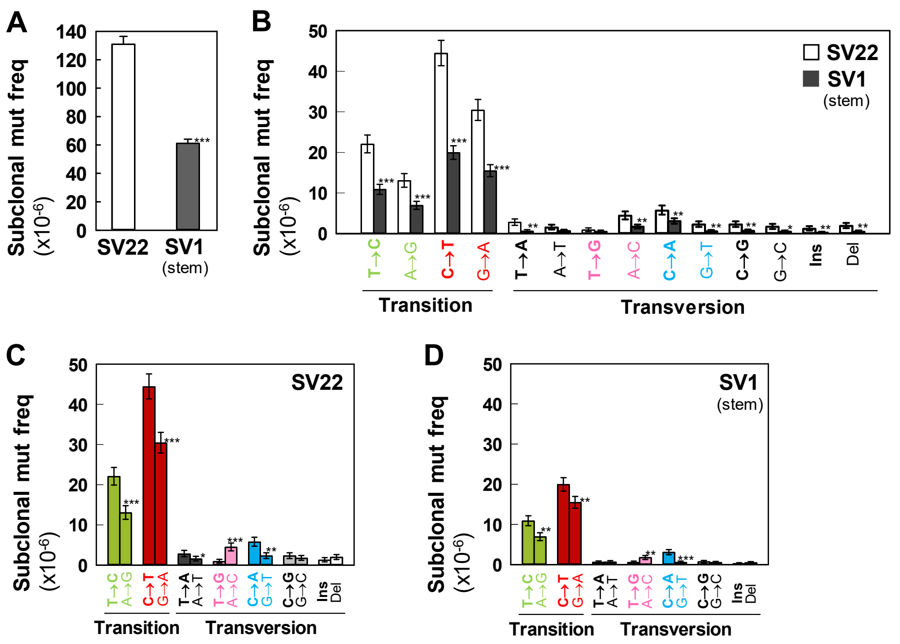


**Figure S2. Frequencies of subclonal mutations in the whole mtDNA.** Overall subclonal mutation frequency (**A**) and frequencies of each type of subclonal mutation (**B-D**) for SV22 (immortalized non-stem) and SV1 (immortalized stem) cells were determined using Duplex Sequencing. Error bars represent the Wilson Score 95% confidence intervals. Significant differences in subclonal mutation frequencies between two groups are indicated (* $p < 0.05$, ** $p < 5 \times 10^{-4}$, and *** $p < 5 \times 10^{-10}$) by the 2-sample test for equality of proportions with continuity correction.

**A**

| | C>A/ G>T | C>G/ G>C | C>T/ G>A | T>A/ A>T | T>C/ A>G | T>G/ A>C |
|---|---|---|---|---|---|---|
| SV22 | 6.0 | 3.0 | 57.0 | 3.3 | 26.7 | 4.0 |
| SV1 (stem) | 5.9 | 1.8 | 57.7 | 2.0 | 29.0 | 3.6 |

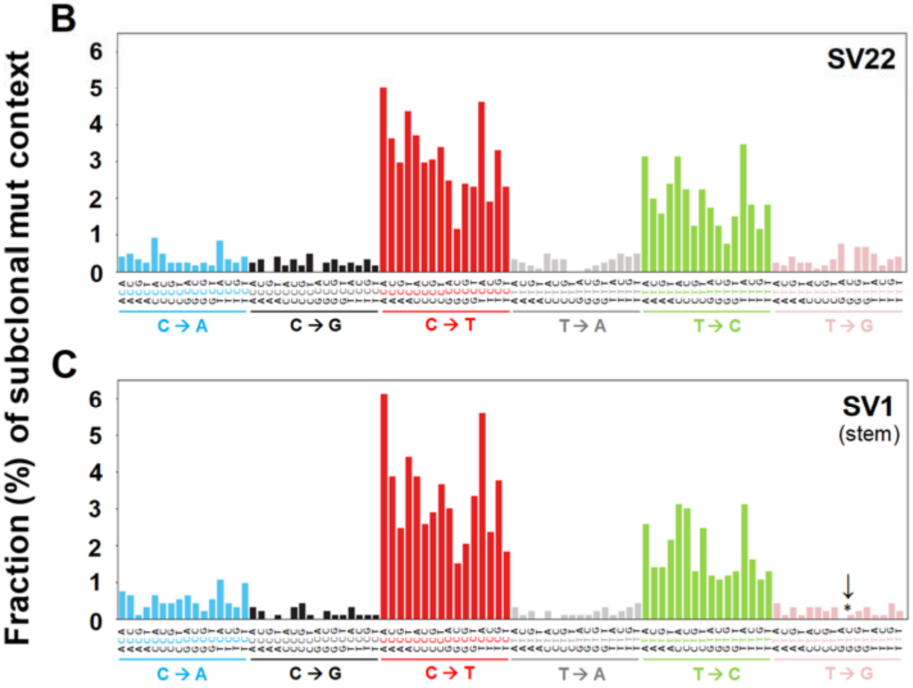


**Figure S3. Types and sequence context spectra of subclonal unique mutations in the whole mtDNA.** Fractions (%) of each type of subclonal unique mutation (**A**) and fractions (%) of subclonal mutation context spectra (**B,C**) for SV22 (immortalized non-stem) and SV1 (immortalized stem) cells were determined using Duplex Sequencing. Trinucleotide contexts (**B,C**) are the mutated base surrounded by all possibilities for its immediate 5' and 3' bases. To keep the graph concise, these point mutation trinucleotides are complemented as necessary to always depict the reference base as the pyrimidine of its pair. The fraction (%) of each specific trinucleotide out of all 96 possible trinucleotide contexts depicts the contribution of each genome sequence context to each point mutation type. Significant differences in fractions (%) of mutation context types between the two groups are indicated (* $p < 0.05$) by the 2-sample test for equality of proportions with continuity correction.

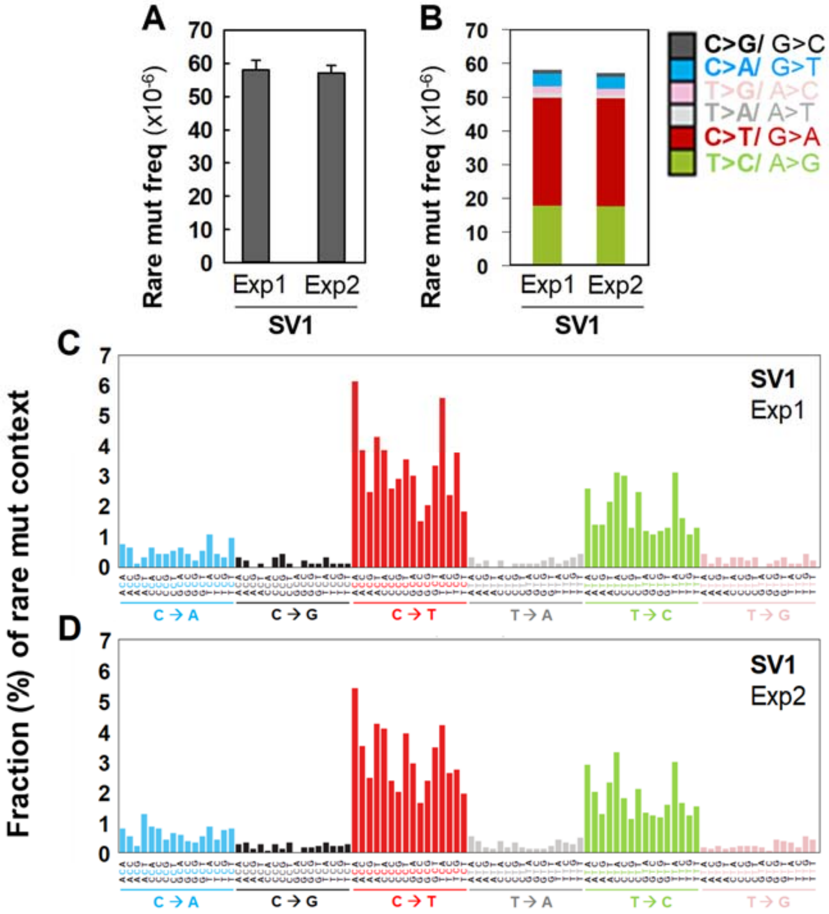


**Figure S4. Reproducibility of DS data: Frequencies and fractions (%) of sequence context spectra of rare mutations in the whole mtDNA.** Data were generated by conducting two independent DNA library experiments (Exp1 and Exp2). Overall rare mutation frequency (**A**), frequencies of rare mutation types (**B**), and fractions (%) of rare unique mutation context spectra (**C,D**) were determined using Duplex Sequencing for immortalized stem cells (SV1). Error bars represent the Wilson Score 95% confidence intervals.

**Table S1.** The number of rare (**A**) and subclonal (**B**) unique mutations in different regions of the whole mtDNA in immortalized human breast epithelial cells (SV22 and SV1). Only genome positions that had a DCS read depth of ≥100 in both samples were considered.

**A.** Rare unique mutations (0 to1%)

| | | **SV22** | **SV1** |
|---|---|---|---|
| Protein coding region | | 727 | 671 |
| Non-coding RNA | mt-RNR1 (rRNA12S) | 171 | 87 |
| | mt-RNR2 (rRNA16S) | 186 | 113 |
| | tRNA | 130 | 78 |
| | Sum | 487 | 278 |
| Control region & overlapped | | 28 | 31 |
| Sum | | 1242 | 980 |

**B.** Subclonal unique mutations (0 to10%)

| | | **SV22** | **SV1** |
|---|---|---|---|
| Protein coding region | | 735 | 677 |
| Non-coding RNA | mt-RNR1 (rRNA12S) | 175 | 87 |
| | mt-RNR2 (rRNA16S) | 187 | 114 |
| | tRNA | 130 | 78 |
| | Sum | 492 | 279 |
| Control region & overlapped | | 29 | 32 |
| Sum | | 1256 | 988 |

Abbreviations used are: SV22, immortalized human breast epithelial cells (HBECs) derived from normal non-stem cells; SV1, immortalized HBECs derived from normal stem cells; DCS, duplex consensus sequence.

**Table S2.** The number of rare unique mutations per 1000 bases in different regions of the whole mtDNA in immortalized human breast epithelial cells (SV22 and SV1). Only genome positions that had a sequence read depth of ≥100 in both samples were considered.

| | | No. of bases on the whole mtDNA | **SV22** | **SV1** |
|---|---|---|---|---|
| Protein coding region | | 11341 | 47 | 44 |
| Non-coding RNA | mt-RNR1 (rRNA12S) | 954 | 179 | 91 |
| | mt-RNR2 (rRNA16S) | 1559 | 119 | 73 |
| | tRNA | 1507 | 86 | 52 |
| | Sum | 4020 | 121 | 69 |

Abbreviations used are: mt, mitochondrial; SV22, immortalized human breast epithelial cells (HBECs) derived from normal non-stem cells; SV1, immortalized HBECs derived from normal stem cells; DCS, duplex consensus sequence.

**Table S3.** Distribution of nonsynonymous unique mutations of subclonal variants within mtDNA coding regions in immortalized human breast epithelial cells (SV22 and SV1). Only genome positions that had a DCS read depth of ≥100 in both samples were considered.

**A.** Rare unique mutations (0 to 1%)

| Cells | Whole mtDNA | Within mtDNA coding regions | | | | Among unique mutations occurring within mtDNA coding regions | |
|---|---|---|---|---|---|---|---|
| | Total sequenced DCS Nts | No. of Syn | No. of Misse | No. of Nonsense | No. of Nonsyn | % Syn | % Nonsyn |
| SV22 | 17562378 | 511 | 207 | 9 | 216 | 70.3 | 29.7 |
| SV1 | 27605473 | 463 | 196 | 12 | 208 | 69.0 | 31.0 |

**B.** Subclonal unique mutations (0 to 10%)

| Cells | Whole mtDNA | Within mtDNA coding regions | | | | Among unique mutations occurring within mtDNA coding regions | |
|---|---|---|---|---|---|---|---|
| | Total sequenced DCS Nts | No. of Syn. | No. of Misse | No. of Nonsense | No. of Nonsyn | % Syn | % Nonsyn |
| SV22 | 17562378 | 513 | 213 | 9 | 222 | 69.8 | 30.2 |
| SV1 | 27605473 | 463 | 202 | 12 | 214 | 68.4 | 31.6 |

Abbreviations used are: Nts, nucleotides; DCS, duplex consensus sequences; mt, mitochondrial; Nonsyn, nonsynonymous mutation; Misse, missense mutation; Syn, synonymous mutation. SV22, immortalized human breast epithelial cells (HBECs) derived from normal non-stem cells; SV1, immortalized HBECs derived from normal stem cells. The numbers of nonsynonymous mutations are sums of missense and nonsense (truncating) mutations.

**Table S4.** Number of unique mutations identified in the whole mtDNA of immortalized human breast epithelial cells (SV22 and SV1) using Duplex Sequencing. Only genome positions that had a DCS read depth of ≥100 in both samples were considered.

| | No. of unique mutations | | |
|---|---|---|---|
| | 0-1% | 0-10% | 0-100% |
| SV22 only [a] | 792 | 797 | 797 |
| SV1 only [b] | 527 | 528 | 528 |
| Both SV22 and SV1 [c] | 447 | 459 | 501 |
| SV22 ↑3X > SV1 [d] | 88 | 92 | 93 |
| SV22 < SV1 ↑3X [e] | 9 | 11 | 11 |

Abbreviations used are:
SV22, immortalized human breast epithelial cells (HBECs) derived from normal non-stem cells; SV1, immortalized HBECs derived from normal stem cells; DCS, duplex consensus sequence.

[a] Mutations found only in SV22 cells but not in SV1 cells
[b] Mutations found only in SV1 cells but not in SV22 cells
[c] Mutations found in both SV22 and SV1 cells
[d] Mutations found in both SV22 and SV1 cells: With being more highly mutated at ≧ 3-fold in SV22 than in SV1 cells
[e] Mutations found in both SV22 and SV1 cells: With being more highly mutated at ≧ 3-fold in SV1 than in SV22 cells

**Table S5.** Mutations identified using Duplex Sequencing in the whole mtDNA of immortalized human breast epithelial cells (SV22 and SV1).

Abbreviations used are: mt, mitochondrial; Nonsyn, nonsynonymous mutation; Syn, synonymous mutation. SV22, immortalized human breast epithelial cells (HBECs) derived from normal non-stem cells; SV1, immortalized HBECs derived from normal stem cells.

**A.** Subclonal variants (excluding singlets) found only in SV22 cells, but not in SV1 cells
Among the mutations identified, variants that have mutated at least twice in each position are listed below.

| Mt Gene | Mutation type | Amino acid change | | SV22 allele frequency (%) | g-score | mitoTIP predicted score |
|---|---|---|---|---|---|---|
| Control region | C>A | . | Non-coding | 0.20 | . | . |
| Control region | A>G | . | Non-coding | 0.36 | . | . |
| Control region | A>G | . | Non-coding | 2.06 | . | . |
| Control region | C>T | . | Non-coding | 0.16 | . | . |
| MT-RNR1 | G>A | . | ncRNA | 0.17 | . | . |
| MT-RNR1 | C>G | . | ncRNA | 0.19 | . | . |
| MT-RNR1 | G>C | . | ncRNA | 0.19 | . | . |
| MT-RNR1 | T>C | . | ncRNA | 0.19 | . | . |
| MT-RNR1 | G>A | . | ncRNA | 0.20 | . | . |
| MT-RNR1 | A>G | . | ncRNA | 0.20 | . | . |
| MT-RNR1 | C>T | . | ncRNA | 0.20 | . | . |
| MT-RNR1 | T>C | . | ncRNA | 0.20 | . | . |
| MT-RNR1 | C>T | . | ncRNA | 0.20 | . | . |
| MT-RNR1 | G>A | . | ncRNA | 0.20 | . | . |
| MT-RNR1 | A>G | . | ncRNA | 0.21 | . | . |
| MT-RNR1 | G>A | . | ncRNA | 0.21 | . | . |
| MT-RNR1 | C>A | . | ncRNA | 0.22 | . | . |
| MT-RNR1 | C>A | . | ncRNA | 0.23 | . | . |
| MT-RNR1 | A>G | . | ncRNA | 0.25 | . | . |
| MT-RNR1 | G>A | . | ncRNA | 0.27 | . | . |
| MT-RNR1 | A>C | . | ncRNA | 0.27 | . | . |
| MT-RNR1 | T>G | . | ncRNA | 0.29 | . | . |
| MT-RNR1 | C>T | . | ncRNA | 0.29 | . | . |
| MT-RNR1 | G>T | . | ncRNA | 0.31 | . | . |
| MT-RNR1 | C>T | . | ncRNA | 0.33 | . | . |
| MT-RNR1 | C>T | . | ncRNA | 0.34 | . | . |
| MT-RNR1 | C>T | . | ncRNA | 0.34 | . | . |
| MT-RNR1 | C>T | . | ncRNA | 0.35 | . | . |
| MT-RNR1 | T>C | . | ncRNA | 0.35 | . | . |
| MT-RNR1 | T>C | . | ncRNA | 0.37 | . | . |
| MT-RNR1 | C>T | . | ncRNA | 0.38 | . | . |
| MT-RNR1 | C>T | . | ncRNA | 0.39 | . | . |
| MT-RNR1 | C>G | . | ncRNA | 0.47 | . | . |
| MT-RNR1 | G>A | . | ncRNA | 0.51 | . | . |
| MT-RNR1 | G>A | . | ncRNA | 0.55 | . | . |

| | | | | | | |
|---|---|---|---|---|---|---|
| MT-RNR1 | G>A | . | ncRNA | 0.55 | . | . |
| MT-RNR1 | C>T | . | ncRNA | 0.70 | . | . |
| MT-RNR1 | C>T | . | ncRNA | 0.73 | . | . |
| MT-RNR1 | C>A | . | ncRNA | 1.12 | . | . |
| MT-TV | A>G | . | ncRNA | 0.18 | . | 16.7 |
| MT-TV | T>C | . | ncRNA | 0.20 | . | 17.0 |
| MT-RNR2 | C>T | . | ncRNA | 0.15 | . | . |
| MT-RNR2 | T>C | . | ncRNA | 0.16 | . | . |
| MT-RNR2 | T>A | . | ncRNA | 0.16 | . | . |
| MT-RNR2 | A>T | . | ncRNA | 0.18 | . | . |
| MT-RNR2 | A>C | . | ncRNA | 0.18 | . | . |
| MT-RNR2 | T>A | . | ncRNA | 0.18 | . | . |
| MT-RNR2 | G>T | . | ncRNA | 0.18 | . | . |
| MT-RNR2 | A>C | . | ncRNA | 0.18 | . | . |
| MT-RNR2 | C>T | . | ncRNA | 0.19 | . | . |
| MT-RNR2 | A>G | . | ncRNA | 0.19 | . | . |
| MT-RNR2 | G>A | . | ncRNA | 0.19 | . | . |
| MT-RNR2 | C>T | . | ncRNA | 0.20 | . | . |
| MT-RNR2 | G>A | . | ncRNA | 0.20 | . | . |
| MT-RNR2 | T>C | . | ncRNA | 0.21 | . | . |
| MT-RNR2 | G>A | . | ncRNA | 0.23 | . | . |
| MT-RNR2 | T>C | . | ncRNA | 0.23 | . | . |
| MT-RNR2 | A>G | . | ncRNA | 0.23 | . | . |
| MT-RNR2 | G>A | . | ncRNA | 0.27 | . | . |
| MT-RNR2 | G>A | . | ncRNA | 0.27 | . | . |
| MT-RNR2 | G>A | . | ncRNA | 0.27 | . | . |
| MT-RNR2 | T>C | . | ncRNA | 0.27 | . | . |
| MT-RNR2 | T>C | . | ncRNA | 0.27 | . | . |
| MT-RNR2 | C>T | . | ncRNA | 0.28 | . | . |
| MT-RNR2 | G>A | . | ncRNA | 0.28 | . | . |
| MT-RNR2 | C>T | . | ncRNA | 0.32 | . | . |
| MT-RNR2 | T>C | . | ncRNA | 0.33 | . | . |
| MT-RNR2 | C>T | . | ncRNA | 0.35 | . | . |
| MT-RNR2 | T>C | . | ncRNA | 0.36 | . | . |
| MT-RNR2 | G>A | . | ncRNA | 0.36 | . | . |
| MT-RNR2 | A>G | . | ncRNA | 0.38 | . | . |
| MT-RNR2 | G>A | . | ncRNA | 0.39 | . | . |
| MT-RNR2 | C>T | . | ncRNA | 0.45 | . | . |
| MT-RNR2 | G>A | . | ncRNA | 0.48 | . | . |
| MT-RNR2 | C>T | . | ncRNA | 0.54 | . | . |
| MT-RNR2 | C>T | . | ncRNA | 0.54 | . | . |
| MT-RNR2 | T>C | . | ncRNA | 0.58 | . | . |
| MT-RNR2 | C>T | . | ncRNA | 0.59 | . | . |
| MT-RNR2 | C>T | . | ncRNA | 0.69 | . | . |
| MT-ND1 | A>C | Missense | T87P | 0.17 | 0.22 | . |

| | | | | | | |
|---|---|---|---|---|---|---|
| MT-ND2 | C>T | Syn | P138P | 0.27 | . | . |
| MT-ND2 | G>A | Missense | G188E | 0.44 | 0.87 | . |
| MT-ND2 | C>T | Missense | T226M | 0.17 | 0.25 | . |
| MT-ND2 | C>T | Syn | T258T | 0.28 | . | . |
| MT-ND2 | G>A | Syn | Q316Q | 1.17 | . | . |
| MT-ND2 | C>T | Syn | T333T | 0.88 | . | . |
| MT-TW | C>T | . | ncRNA | 0.21 | . | 3.1 |
| MT-TW | A>G | . | ncRNA | 0.21 | . | 1.2 |
| MT-TW | C>T | . | ncRNA | 0.45 | . | 15.5 |
| MT-TA | T>C | . | ncRNA | 0.20 | . | 4.4 |
| MT-TA | T>C | . | ncRNA | 0.20 | . | 16.7 |
| MT-TA | C>T | . | ncRNA | 0.20 | . | 0.9 |
| MT-TA | C>A | . | ncRNA | 0.20 | . | 4.7 |
| MT-COX1 | C>T | Syn | Y54Y | 0.23 | . | . |
| MT-COX1 | A>G | Syn | W81W | 0.14 | . | . |
| MT-COX1 | A>G | Syn | W126W | 0.14 | . | . |
| MT-COX1 | C>T | Syn | H290H | 0.15 | . | . |
| MT-COX1 | T>C | Syn | F293F | 0.17 | . | . |
| MT-COX1 | C>T | Syn | D298D | 0.16 | . | . |
| MT-COX1 | C>T | Syn | I312I | 0.16 | . | . |
| MT-COX1 | C>T | Syn | P315P | 0.17 | . | . |
| MT-COX1 | C>A | Nonsense | S362X | 0.17 | . | . |
| MT-COX1 | C>T | Syn | H376H | 0.22 | . | . |
| MT-COX1 | T>G | Syn | A385A | 0.17 | . | . |
| MT-COX1 | C>T | Syn | Y403Y | 0.38 | . | . |
| MT-COX1 | T>C | Syn | T424T | 0.43 | . | . |
| MT-COX1 | C>T | Syn | I419I | 0.55 | . | . |
| MT-COX2 | C>A | Missense | F36L | 1.66 | 0.12 | . |
| MT-COX2 | G>A | Missense | V38I | 0.36 | 0.18 | . |
| MT-COX2 | C>T | Missense | L84F | 0.20 | 0.65 | . |
| MT-COX2 | C>T | Missense | L144F | 3.32 | 0.68 | . |
| MT-COX2 | C>T | Missense | T226I | 0.80 | 0.12 | . |
| MT-TK | G>A | . | ncRNA | 0.41 | . | 17.5 |
| MT-ATP6 | T>C | Syn | Y36Y | 0.21 | . | . |
| MT-ATP6 | C>T | Syn | I43I | 0.19 | . | . |
| MT-ATP6 | C>T | Missense | R66W | 0.56 | 0.18 | . |
| MT-ATP6 | G>A | Syn | L156L | 0.27 | . | . |
| MT-ATP8 | T>C | Syn | L29L | 0.19 | . | . |
| MT-COX3 | G>A | Syn | M33M | 0.18 | . | . |
| MT-COX3 | C>T | Syn | H115H | 0.48 | . | . |
| MT-ND3 | G>T | Nonsense | E105X | 0.77 | . | . |
| MT-ND4 | C>T | Missense | S101F | 0.16 | 0.11 | . |
| MT-ND4 | A>C | Syn | L102L | 0.24 | . | . |
| MT-ND4 | T>A | Missense | I104N | 0.16 | 0.44 | . |
| MT-ND4 | C>T | Syn | I116I | 0.16 | . | . |

| | | | | | | |
|---|---|---|---|---|---|---|
| MT-ND4 | T>C | Syn | F118F | 0.16 | . | . |
| MT-ND4 | T>C | Syn | Y119Y | 0.17 | . | . |
| MT-ND4 | G>A | Missense | A287T | 0.84 | 0.32 | . |
| MT-ND4 | G>A | Syn | L376L | 0.19 | . | . |
| MT-ND4 | G>T | Syn | L376L | 0.66 | . | . |
| MT-ND4 | G>A | Syn | T385T | 0.17 | . | . |
| MT-ND4 | A>G | Syn | S418S | 0.19 | . | . |
| MT-ND4 | C>G | Missense | T420S | 0.18 | 0.29 | . |
| MT-TS2 | T>C | . | ncRNA | 0.18 | . | 2.9 |
| MT-ND5 | T>A | Nonsense | Y35X | 0.19 | . | . |
| MT-ND5 | G>A | Missense | V40I | 0.27 | 0.06 | . |
| MT-ND5 | C>T | Syn | Y84Y | 0.17 | . | . |
| MT-ND5 | C>T | Missense | L216F | 0.25 | 0.21 | . |
| MT-ND5 | C>T | Syn | L229L | 0.17 | . | . |
| MT-ND5 | C>T | Syn | L266L | 0.21 | . | . |
| MT-ND5 | G>A | Missense | S270N | 0.18 | 0.04 | . |
| MT-ND5 | A>C | Syn | L276L | 0.28 | . | . |
| MT-ND5 | T>C | Syn | L280L | 0.45 | . | . |
| MT-ND5 | T>C | Missense | I283T | 0.19 | 0.36 | . |
| MT-ND5 | T>C | Syn | T285T | 0.29 | . | . |
| MT-ND5 | T>C | Syn | I566I | 0.18 | . | . |
| MT-ND5 | A>G | Missense | T573A | 0.18 | 0.04 | . |
| MT-ND5 | C>T | Syn | T579T | 0.16 | . | . |
| MT-ND5 | A>C | Missense | I596L | 0.18 | 0.07 | . |
| MT-ND5 | T>C | Syn | I584I | 0.21 | . | . |
| MT-ND6 | A>T | Missense | F46Y | 0.29 | 0.50 | . |
| MT-ND6 | C>T | Syn | M51M | 0.20 | . | . |
| MT-ND6 | T>C | Syn | M54M | 0.40 | . | . |
| MT-ND6 | T>A | Syn | G62G | 0.21 | . | . |
| MT-ND6 | C>T | Syn | M63M | 0.38 | . | . |
| MT-ND6 | A>G | Syn | V65V | 0.18 | . | . |
| MT-ND6 | G>A | Syn | V66V | 0.48 | . | . |
| MT-ND6 | A>G | Syn | Y69Y | 0.20 | . | . |
| MT-ND6 | C>T | Syn | A72A | 0.19 | . | . |
| MT-ND6 | C>T | Syn | M73M | 0.29 | . | . |
| MT-ND6 | C>T | Syn | V90V | 0.22 | . | . |
| MT-ND6 | T>C | Syn | K107K | 0.20 | . | . |
| MT-ND6 | A>T | Syn | V114V | 0.19 | . | . |
| MT-ND6 | C>T | Missense | V114I | 0.20 | 0.22 | . |
| MT-ND6 | G>A | Syn | N117N | 0.30 | . | . |
| MT-ND6 | T>C | Syn | G129G | 0.17 | . | . |
| MT-ND6 | C>G | Missense | G133A | 0.16 | 0.79 | . |
| MT-ND6 | C>T | Syn | G133G | 0.16 | . | . |
| MT-ND6 | A>G | Missense | S132P | 0.24 | 0.67 | . |
| MT-ND6 | T>C | Syn | V121V | 0.29 | . | . |

| | | | | | | |
|---|---|---|---|---|---|---|
| MT-CYTB | C>A | Missense | H16Q | 0.15 | 0.20 | . |
| MT-CYTB | T>C | Missense | F18S | 0.30 | 0.39 | . |
| MT-CYTB | G>A | Missense | A29T | 0.16 | 0.09 | . |
| MT-CYTB | T>C | Missense | F33L | 0.16 | 0.30 | . |
| MT-CYTB | C>T | Missense | S110L | 0.17 | 0.14 | . |
| MT-CYTB | T>C | Missense | S110P | 0.17 | 0.43 | . |
| MT-CYTB | T>A | Nonsense | S172X | 0.15 | . | . |
| MT-CYTB | T>C | Syn | F181F | 0.16 | . | . |
| MT-CYTB | C>T | Syn | Y224Y | 0.84 | . | . |
| MT-CYTB | A>G | Syn | L250L | 0.18 | . | . |
| MT-CYTB | G>A | Missense | G251D | 0.18 | 0.36 | . |
| MT-CYTB | T>C | Syn | L262L | 0.28 | . | . |
| MT-CYTB | G>A | Missense | R318H | 0.28 | 0.08 | . |
| MT-CYTB | C>T | Syn | L320L | 0.29 | . | . |
| MT-CYTB | A>C | Syn | S323S | 0.18 | . | . |
| MT-CYTB | C>T | Missense | S323L | 0.18 | 0.03 | . |
| MT-CYTB | T>A | Syn | L324L | 0.18 | . | . |
| MT-CYTB | C>T | Missense | L327F | 0.19 | 0.09 | . |
| MT-CYTB | G>A | Missense | A330T | 0.20 | 0.09 | . |
| MT-TT | G>A | . | ncRNA | 0.19 | . | 2.4 |

**B.** Subclonal variants (excluding singlets) found only in SV1 cells, but not in SV22 cells
Among the mutations identified, variants that have mutated at least twice in each position are listed below.

| Mt Gene | Mutation type | Amino acid change | | SV1 allele frequency (%) | g-score | mitoTIP predicted score |
|---|---|---|---|---|---|---|
| Control region | G>C | . | Non-coding | 1.69 | . | . |
| Control region | C>T | . | Non-coding | 0.47 | . | . |
| MT-RNR1 | T>C | . | ncRNA | 0.09 | . | . |
| MT-RNR1 | G>A | . | ncRNA | 0.09 | . | . |
| MT-RNR1 | C>T | . | ncRNA | 0.08 | . | . |
| MT-RNR2 | A>G | . | ncRNA | 0.11 | . | . |
| MT-RNR2 | G>A | . | ncRNA | 0.11 | . | . |
| MT-RNR2 | G>C | . | ncRNA | 0.13 | . | . |
| MT-RNR2 | T>C | . | ncRNA | 0.11 | . | . |
| MT-RNR2 | T>C | . | ncRNA | 0.10 | . | . |
| MT-RNR2 | C>T | . | ncRNA | 0.15 | . | . |
| MT-RNR2 | G>A | . | ncRNA | 0.16 | . | . |
| MT-RNR2 | C>T | . | ncRNA | 0.10 | . | . |
| MT-RNR2 | A>G | . | ncRNA | 0.10 | . | . |
| MT-RNR2 | C>T | . | ncRNA | 0.19 | . | . |
| MT-ND1 | T>C | Missense | F56S | 0.14 | 0.69 | . |
| MT-ND1 | A>C | Missense | T57P | 0.23 | 0.22 | . |
| MT-ND1 | G>C | Missense | E59D | 0.16 | 0.21 | . |
| MT-ND1 | T>A | Missense | L61Q | 0.61 | 0.47 | . |

| | | | | | | |
|---|---|---|---|---|---|---|
| MT-ND1 | A>C | Missense | K62N | 0.36 | 0.16 | . |
| MT-ND1 | G>A | Syn | L176L | 0.10 | . | . |
| MT-ND1 | T>C | Missense | Y304H | 0.36 | 0.09 | . |
| MT-TN | A>C | . | ncRNA | 0.22 | . | 8.3 |
| MT-COX1 | G>A | Syn | E40E | 0.16 | . | . |
| MT-COX1 | A>G | Syn | V70V | 0.13 | . | . |
| MT-COX1 | C>T | Syn | H151H | 0.12 | . | . |
| MT-COX1 | G>A | Missense | V155I | 0.28 | 0.06 | . |
| MT-COX1 | A>C | Missense | K172N | 0.28 | 0.39 | . |
| MT-COX1 | C>A | Syn | G250G | 0.08 | . | . |
| MT-COX1 | C>T | Syn | Y260Y | 0.08 | . | . |
| MT-COX1 | T>C | Missense | S262P | 0.08 | 0.59 | . |
| MT-COX1 | A>G | Syn | K265K | 0.07 | . | . |
| MT-COX1 | T>C | Syn | G272G | 0.07 | . | . |
| MT-COX1 | G>A | Syn | G284G | 0.11 | . | . |
| MT-COX1 | A>T | Syn | A303A | 0.08 | . | . |
| MT-COX1 | G>A | Missense | A313T | 0.08 | 0.47 | . |
| MT-COX1 | C>T | Syn | S322S | 0.08 | . | . |
| MT-COX1 | A>G | Syn | W323W | 0.08 | . | . |
| MT-COX1 | A>G | Syn | A337A | 0.08 | . | . |
| MT-COX1 | A>G | Syn | L342L | 0.12 | . | . |
| MT-COX1 | A>G | Syn | V350V | 0.13 | . | . |
| MT-COX2 | C>T | Syn | Y105Y | 0.09 | . | . |
| MT-COX2 | C>T | Missense | P125L | 0.16 | 0.15 | . |
| MT-COX2 | G>A | Syn | L160L | 0.23 | . | . |
| MT-COX2 | G>A | Syn | P189P | 0.19 | . | . |
| MT-COX2 | T>C | Syn | G194G | 0.19 | . | . |
| MT-ATP8 | C>A | Missense | P53Q | 0.17 | 0.16 | . |
| MT-COX3 | T>C | Syn | L137L | 0.09 | . | . |
| MT-COX3 | T>C | Syn | A147A | 0.08 | . | . |
| MT-COX3 | A>C | Syn | V142V | 0.09 | . | . |
| MT-COX3 | C>T | Syn | T145T | 0.09 | . | . |
| MT-COX3 | G>A | Syn | L169L | 0.12 | . | . |
| MT-COX3 | C>G | Missense | L171V | 0.12 | 0.21 | . |
| MT-COX3 | C>T | Syn | L171L | 0.12 | . | . |
| MT-COX3 | T>C | Syn | Y172Y | 0.12 | . | . |
| MT-COX3 | C>T | Syn | Y181Y | 0.13 | . | . |
| MT-ND3 | A>C | Missense | T35P | 0.12 | 0.49 | . |
| MT-ND3 | A>C | Missense | T61P | 0.10 | 0.86 | . |
| MT-ND3 | C>T | Syn | L86L | 0.09 | . | . |
| MT-ND3 | T>C | Syn | V88V | 0.09 | . | . |
| MT-ND3 | G>A | Syn | M89M | 0.09 | . | . |
| MT-ND3 | T>C | Missense | M89T | 0.09 | 0.07 | . |
| MT-ND3 | A>G | Syn | S90S | 0.09 | . | . |
| MT-ND3 | T>C | Syn | Y104Y | 0.09 | . | . |

| | | | | | | |
|---|---|---|---|---|---|---|
| MT-ND3 | G>A | Syn | E105E | 0.09 | . | . |
| MT-ND3 | C>T | Syn | L107L | 0.09 | . | . |
| MT-ND3 | C>T | Nonsense | Q108X | 0.09 | . | . |
| MT-ND3 | A>G | Syn | G110G | 0.13 | . | . |
| MT-TR | T>C | . | ncRNA | 0.17 | . | 17.8 |
| MT-ND4L | G>A | Syn | M1M | 0.09 | . | . |
| MT-ND4L | T>C | Syn | A44A | 0.14 | . | . |
| MT-ND4 | T>C | Syn | I9I | 0.19 | . | . |
| MT-ND4 | G>A | Syn | L14L | 0.15 | . | . |
| MT-ND4 | C>T | Missense | A131V | 0.10 | 0.50 | . |
| MT-ND4 | T>A | Syn | A198A | 0.11 | . | . |
| MT-ND4 | T>C | Syn | F203F | 0.11 | . | . |
| MT-ND4 | G>A | Syn | K206K | 0.17 | . | . |
| MT-ND4 | A>G | Syn | M207M | 0.12 | . | . |
| MT-ND4 | T>C | Syn | P208P | 0.12 | . | . |
| MT-ND4 | T>C | Syn | L209L | 0.12 | . | . |
| MT-ND4 | G>A | Nonsense | G211X | 0.12 | . | . |
| MT-ND4 | T>C | Syn | L214L | 0.24 | . | . |
| MT-ND4 | A>G | Syn | W215W | 0.12 | . | . |
| MT-ND4 | A>G | Syn | V234V | 0.14 | . | . |
| MT-ND4 | C>T | Syn | G239G | 0.15 | . | . |
| MT-ND4 | T>C | Syn | G242G | 0.15 | . | . |
| MT-ND4 | A>G | Syn | M244M | 0.15 | . | . |
| MT-ND4 | A>C | Syn | T247T | 0.16 | . | . |
| MT-ND4 | T>C | Syn | T337T | 0.12 | . | . |
| MT-ND4 | C>T | Syn | N425N | 0.41 | . | . |
| MT-ND4 | G>A | Syn | M437M | 0.11 | . | . |
| MT-ND4 | T>C | Syn | I444I | 0.11 | . | . |
| MT-ND4 | C>T | Syn | L449L | 0.11 | . | . |
| MT-ND4 | C>T | Syn | D452D | 0.11 | . | . |
| MT-ND4 | T>C | Syn | S459S | 0.12 | . | . |
| MT-ND4 | C>T | Syn | P451P | 0.11 | . | . |
| MT-ND5 | C>T | Missense | H4Y | 0.12 | 0.12 | . |
| MT-ND5 | A>C | Missense | T8P | 0.18 | 0.28 | . |
| MT-ND5 | A>G | Missense | T11A | 0.18 | 0.11 | . |
| MT-ND5 | C>T | Syn | L15L | 0.18 | . | . |
| MT-ND5 | C>T | Syn | P18P | 0.18 | . | . |
| MT-ND5 | G>A | Missense | S47N | 0.51 | 0.28 | . |
| MT-ND5 | C>T | Syn | I123I | 0.11 | . | . |
| MT-ND5 | C>A | Syn | P240P | 0.27 | . | . |
| MT-ND5 | C>T | Syn | V243V | 0.08 | . | . |
| MT-ND5 | C>T | Syn | A245A | 0.08 | . | . |
| MT-ND5 | C>T | Missense | A245V | 0.08 | 0.33 | . |
| MT-ND5 | G>A | Missense | S377N | 0.12 | 0.09 | . |
| MT-ND5 | A>G | Syn | L380L | 0.13 | . | . |

| | | | | | | |
|---|---|---|---|---|---|---|
| MT-ND5 | G>A | Missense | A399T | 0.14 | 0.16 | . |
| MT-ND5 | A>C | Syn | T404T | 0.13 | . | . |
| MT-ND5 | T>C | Syn | A415A | 0.12 | . | . |
| MT-ND5 | C>T | Missense | S473F | 0.21 | 0.20 | . |
| MT-CYTB | G>A | Syn | M53M | 0.16 | . | . |
| MT-CYTB | C>T | Syn | A59A | 0.15 | . | . |
| MT-CYTB | A>G | Syn | S65S | 0.10 | . | . |
| MT-CYTB | C>T | Syn | H68H | 0.14 | . | . |
| MT-CYTB | G>A | Syn | P134P | 0.15 | . | . |
| MT-CYTB | A>C | Syn | S139S | 0.23 | . | . |
| MT-CYTB | C>A | Missense | I211M | 0.12 | 0.12 | . |
| MT-CYTB | G>A | Missense | S344N | 0.17 | 0.26 | . |
| MT-CYTB | C>T | Syn | T348T | 0.17 | . | . |

**C.** Subclonal variants found in both SV22 and SV1 cells: With being more highly mutated at ≧ 3-fold in SV22 than in SV1 cells

| Mt Gene | DNA mutation | Amino acid change | | SV22 allele frequency (%) | SV1 allele frequency (%) | Fold Change (SV22/SV1) | g-score | mitoTIP predicted score |
|---|---|---|---|---|---|---|---|---|
| Control region | G>C | . | Non-coding | 0.29 | 0.04 | 6.62 | . | . |
| Control region | C>G | . | Non-coding | 0.19 | 0.04 | 4.85 | . | . |
| MT-RNR1 | A>G | . | ncRNA | 0.26 | 0.06 | 4.42 | . | . |
| MT-RNR1 | T>C | . | ncRNA | 0.31 | 0.05 | 5.71 | . | . |
| MT-RNR1 | A>G | . | ncRNA | 0.30 | 0.05 | 6.01 | . | . |
| MT-RNR1 | T>A | . | ncRNA | 0.52 | 0.06 | 9.30 | . | . |
| MT-RNR1 | C>T | . | ncRNA | 0.21 | 0.07 | 3.21 | . | . |
| MT-RNR1 | G>A | . | ncRNA | 0.75 | 0.19 | 4.00 | . | . |
| MT-RNR1 | G>A | . | ncRNA | 0.45 | 0.10 | 4.41 | . | . |
| MT-RNR1 | C>G | . | ncRNA | 0.49 | 0.07 | 6.75 | . | . |
| MT-RNR1 | A>G | . | ncRNA | 0.21 | 0.07 | 3.10 | . | . |
| MT-RNR1 | C>T | . | ncRNA | 0.43 | 0.12 | 3.46 | . | . |
| MT-RNR1 | G>A | . | ncRNA | 0.87 | 0.17 | 5.08 | . | . |
| MT-RNR1 | C>A | . | ncRNA | 0.89 | 0.17 | 5.26 | . | . |
| MT-RNR1 | C>T | . | ncRNA | 1.20 | 0.20 | 6.04 | . | . |
| MT-RNR1 | G>A | . | ncRNA | 0.19 | 0.05 | 3.95 | . | . |
| MT-RNR1 | C>T | . | ncRNA | 1.18 | 0.10 | 11.41 | . | . |
| MT-RNR1 | G>A | . | ncRNA | 0.42 | 0.05 | 8.04 | . | . |
| MT-RNR1 | G>A | . | ncRNA | 0.89 | 0.15 | 5.84 | . | . |
| MT-RNR1 | A>T | . | ncRNA | 0.21 | 0.05 | 4.28 | . | . |
| MT-RNR1 | T>C | . | ncRNA | 0.46 | 0.05 | 9.71 | . | . |
| MT-RNR1 | G>A | . | ncRNA | 0.23 | 0.04 | 5.12 | . | . |
| MT-RNR1 | G>T | . | ncRNA | 0.46 | 0.04 | 10.24 | . | . |

| | | | | | | | | |
|---|---|---|---|---|---|---|---|---|
| MT-RNR1 | T>C | . | ncRNA | 0.54 | 0.05 | 11.84 | . | . |
| MT-RNR1 | C>T | . | ncRNA | 0.40 | 0.04 | 9.35 | . | . |
| MT-RNR1 | T>C | . | ncRNA | 0.22 | 0.04 | 5.05 | . | . |
| MT-RNR1 | G>C | . | ncRNA | 0.22 | 0.05 | 4.76 | . | . |
| MT-RNR1 | T>A | . | ncRNA | 0.21 | 0.05 | 4.32 | . | . |
| MT-RNR1 | G>A | . | ncRNA | 1.26 | 0.42 | 3.02 | . | . |
| MT-RNR2 | A>G | . | ncRNA | 0.34 | 0.10 | 3.47 | . | . |
| MT-RNR2 | G>A | . | ncRNA | 0.45 | 0.05 | 8.32 | . | . |
| MT-RNR2 | C>T | . | ncRNA | 0.27 | 0.06 | 4.11 | . | . |
| MT-RNR2 | G>A | . | ncRNA | 0.30 | 0.06 | 4.89 | . | . |
| MT-RNR2 | C>T | . | ncRNA | 0.28 | 0.07 | 3.91 | . | . |
| MT-RNR2 | T>C | . | ncRNA | 0.38 | 0.07 | 5.25 | . | . |
| MT-RNR2 | C>T | . | ncRNA | 0.30 | 0.07 | 4.04 | . | . |
| MT-RNR2 | T>C | . | ncRNA | 0.30 | 0.07 | 4.15 | . | . |
| MT-RNR2 | T>C | . | ncRNA | 0.45 | 0.09 | 4.90 | . | . |
| MT-RNR2 | C>T | . | ncRNA | 0.47 | 0.10 | 4.60 | . | . |
| MT-RNR2 | G>A | . | ncRNA | 9.63 | 1.53 | 6.30 | . | . |
| MT-RNR2 | G>A | . | ncRNA | 0.50 | 0.07 | 7.20 | . | . |
| MT-TL1 | G>A | . | ncRNA | 0.32 | 0.05 | 5.82 | . | 2.1 |
| MT-TL1 | T>C | . | ncRNA | 0.37 | 0.05 | 6.97 | . | 2.0 |
| MT-TL1 | C>T | . | ncRNA | 0.31 | 0.05 | 5.76 | . | 1.5 |
| MT-ND2 | A>G | Syn | M163M | 0.30 | 0.05 | 5.84 | . | . |
| MT-ND2 | G>A | Missense | Q174Q | 0.55 | 0.15 | 3.58 | . | . |
| MT-ND2 | T>C | Syn | M191T | 0.21 | 0.04 | 4.93 | 0.22 | . |
| MT-TA | C>T | . | ncRNA | 0.46 | 0.04 | 11.78 | . | 6.3 |
| MT-TA | C>T | . | ncRNA | 0.41 | 0.04 | 9.98 | . | 12.3 |
| MT-TA | G>A | . | ncRNA | 0.45 | 0.08 | 5.79 | . | 12.0 |
| MT-TN | C>T | . | ncRNA | 0.36 | 0.04 | 9.15 | . | 2.4 |
| MT-TN | G>A | . | ncRNA | 0.74 | 0.21 | 3.59 | . | 4.0 |
| MT-TN | C>T | . | ncRNA | 0.37 | 0.08 | 4.58 | . | 10.4 |
| MT-TN | C>G | . | ncRNA | 0.17 | 0.04 | 4.44 | . | 18.1 |
| MT-TC | C>T | . | ncRNA | 0.18 | 0.04 | 4.88 | . | 13.6 |
| MT-TC | T>A | . | ncRNA | 0.36 | 0.08 | 4.75 | . | 1.2 |
| MT-TC | T>C | . | ncRNA | 0.17 | 0.04 | 4.61 | . | 2.5 |
| MT-TC | C>T | . | ncRNA | 0.27 | 0.08 | 3.49 | . | -1.7 |
| MT-COX1 | C>T | Syn | L183L | 0.20 | 0.04 | 4.71 | . | . |
| MT-COX1 | C>T | Syn | F238F | 0.25 | 0.07 | 3.43 | . | . |
| MT-COX1 | T>C | Syn | G239G | 0.17 | 0.04 | 4.50 | . | . |
| MT-COX1 | T>C | Syn | P241P | 0.17 | 0.04 | 4.69 | . | . |
| MT-COX1 | T>C | Syn | V243V | 0.17 | 0.04 | 4.61 | . | . |

| | | | | | | | | |
|---|---|---|---|---|---|---|---|---|
| MT-COX1 | C>T | Syn | F251F | 0.19 | 0.04 | 4.82 | . | . |
| MT-COX1 | A>G | Syn | W275W | 0.16 | 0.04 | 4.45 | . | . |
| MT-COX1 | G>A | Syn | M277M | 0.24 | 0.04 | 6.76 | . | . |
| MT-COX1 | A>C | Syn | A289A | 0.23 | 0.04 | 6.21 | . | . |
| MT-COX1 | T>C | Syn | Y304Y | 0.25 | 0.04 | 6.03 | . | . |
| MT-COX1 | C>T | Syn | L363L | 0.33 | 0.11 | 3.12 | . | . |
| MT-COX1 | G>A | Syn | T370T | 0.40 | 0.11 | 3.64 | . | . |
| MT-COX1 | T>C | Syn | V373V | 0.39 | 0.11 | 3.50 | . | . |
| MT-COX1 | A>G | Syn | L399L | 0.31 | 0.06 | 5.42 | . | . |
| MT-COX1 | C>T | Syn | A410A | 0.20 | 0.06 | 3.15 | . | . |
| MT-COX1 | C>T | Syn | F414F | 0.68 | 0.06 | 11.02 | . | . |
| MT-COX1 | T>C | Syn | N422N | 0.86 | 0.07 | 12.71 | . | . |
| MT-COX1 | C>T | Syn | L431L | 0.37 | 0.07 | 5.23 | . | . |
| MT-COX1 | C>T | Syn | S434S | 0.46 | 0.07 | 6.37 | . | . |
| MT-COX1 | C>T | Syn | N451N | 0.60 | 0.16 | 3.67 | . | . |
| MT-ATP6 | A>G | Missense | M1V | 0.31 | 0.08 | 3.95 | 0.76 | . |
| MT-ATP8 | A>G | Syn | K54K | 0.31 | 0.08 | 3.95 | . | . |
| MT-COX3 | T>C | Missense | M40T | 0.31 | 0.10 | 3.13 | 0.07 | . |
| MT-ND5 | C>T | Syn | Y32Y | 0.18 | 0.05 | 3.27 | . | . |
| MT-ND5 | G>A | Missense | Q116Q | 0.17 | 0.06 | 3.00 | . | . |
| MT-ND5 | C>T | Syn | A225A | 0.25 | 0.04 | 5.89 | . | . |
| MT-ND5 | T>A | Syn | L246Q | 0.19 | 0.04 | 4.58 | 0.60 | . |
| MT-ND6 | T>C | Syn | G68G | 0.39 | 0.07 | 5.95 | . | . |
| MT-ND6 | G>A | Syn | S123S | 0.36 | 0.06 | 6.35 | . | . |
| MT-CYTB | C>T | Syn | L36L | 0.25 | 0.05 | 5.27 | . | . |
| MT-CYTB | G>A | Syn | L41L | 0.19 | 0.05 | 3.72 | . | . |
| MT-CYTB | T>C | Syn | N74N | 0.27 | 0.04 | 6.15 | . | . |
| MT-CYTB | G>C | Syn | G99G | 0.27 | 0.05 | 5.09 | . | . |
| MT-CYTB | A>C | Syn | G105G | 0.18 | 0.06 | 3.12 | . | . |

**D.** Subclonal variants found in both SV22 and SV1 cells: With being more highly mutated at ≥ 3- fold in SV1 than in SV22 cells

| Mt Gene | DNA mutation | Amino acid change | | SV22 allele frequency (%) | SV1 allele frequency (%) | Fold Change (SV1/SV22) | g-score | mitoTIP predicted score |
|---|---|---|---|---|---|---|---|---|
| MT-TY | C>T | . | ncRNA | 0.08 | 0.38 | 4.98 | . | 5.0 |
| MT-COX1 | C>T | Syn | A325A | 0.08 | 0.44 | 5.47 | . | . |
| MT-TD | G>A | . | ncRNA | 0.16 | 0.74 | 4.65 | . | 0.71 |
| MT-COX2 | G>A | Syn | E109E | 0.08 | 0.32 | 3.88 | . | . |
| MT-ATP6 | G>C | Missense | A11P | 0.32 | 1.44 | 4.43 | 0.27 | . |
| MT-ATP8 | G>C | Missense | L64F | 0.32 | 1.44 | 4.43 | 0.12 | . |

| | | | | | | | | |
|---|---|---|---|---|---|---|---|---|
| MT-COX3 | C>T | Syn | A107A | 0.09 | 0.30 | 3.25 | . | . |
| MT-COX3 | T>C | Syn | P108P | 0.10 | 0.31 | 3.15 | . | . |
| MT-COX3 | T>C | Syn | L112L | 0.10 | 0.40 | 4.01 | . | . |
| MT-COX3 | A>G | Syn | G113G | 0.10 | 0.35 | 3.46 | . | . |
| MT-COX3 | G>A | Syn | G114G | 0.10 | 0.39 | 4.03 | . | . |
